# Linking clinical border morphology to cell-scale growth mechanisms in melanoma: a segmentation-to-simulation alignment study

**DOI:** 10.64898/2026.10.26.761769

**Authors:** Chang Liu, Lingling Zu

**Affiliations:** School of Medicine, Nankai University, Tianjin 300071, China; Tianjin Key Laboratory of Lung Cancer Metastasis and Tumor Microenvironment, Tianjin Lung Cancer Institute, Tianjin Medical University General Hospital, Tianjin, China

## Abstract

An irregular border is among the most informative visual cues for melanoma, yet the prevailing quantitative treatment of that irregularity, the box-counting fractal dimension, has rarely been connected to the cell-scale mechanism that produces it. We measured seven boundary descriptors on 1,581 lesions from the 2018 International Skin Imaging Collaboration archive segmented by an automatic thresholding route, retaining the expert ground-truth route as a sensitivity arm, and compared their medians across benign, in situ, and invasive grades. We then built a hybrid cellular automaton and swept four canonical cell-level mechanisms, scoring every simulated boundary with the same measurement code used on the patient images, before testing two mechanisms that lift the empty-neighbour division rule and aligning them to the observed malignant signature with a variance-based sensitivity analysis. The malignant signature is lobulation, not roughness: compactness rises from 1.351 to 1.810, solidity falls from 0.910 to 0.853, and convexity falls from 0.936 to 0.839 from benign to invasive lesions, with areas under the curve near 0.74, while the fractal dimension varies by only 0.024 across the three grades. All four canonical mechanisms, including loss of contact inhibition, converge on near-circular clusters at a distance of 1.147 to 1.166 from the malignant target. A long-range spatial heterogeneity in the growth rate, by contrast, reproduces the observed signature to a distance of 0.187, with the correlation length of the field, not its amplitude, the dominant parameter by total-effect index. A smooth, long-range growth-rate field is therefore sufficient to generate the lobular borders that characterise invasive melanoma, whereas the empty-neighbour assumption common to cellular automata is not. Because the winning field is imposed rather than emergent, the study resolves the alignment between border morphology and cell-scale mechanism while leaving open how such heterogeneity could arise from cell-autonomous dynamics.

**Author Summary:** Melanoma is the deadliest form of skin cancer, and clinicians have long used the raggedness of a lesion’s edge as a warning sign. Computer methods usually summarise that raggedness with a single number, the fractal dimension, which measures how much fine detail a border contains. We tested that habit on 1,581 skin lesions and found it misleading. What distinguishes malignant from benign lesions here is not fine roughness but a handful of large lobes: the outline bulges and dents at a coarse scale rather than fraying at a fine one. We then asked what kind of cell behaviour could build such a border. We grew virtual tumours using simple rules for cell division, movement, and crowding, and we measured their edges with exactly the same code we applied to the patient images. The usual suspects, namely cells that ignore crowding, migrate further, or vary in their growth, all produced smooth and round tumours. Only when we let the growth rate vary smoothly across the tumour did the simulated edge acquire the observed lobes. We could impose that variation but cannot yet explain where it comes from, and we say so. That open question is our next target.

## Introduction

Cutaneous melanoma is the most lethal form of skin cancer: it accounts for only a small fraction of skin cancer cases yet the large majority of skin cancer deaths, and its prognosis is determined in large part by how early the lesion is recognised. Its incidence has risen in many populations and is projected to increase further [1], within a global cancer burden that reached an estimated 20 million new cases in 2022 [2]. Among the visual cues a clinician weighs, the irregularity of the lesion border is one of the most informative, formalised as the border criterion of the ABCD screening rule alongside asymmetry [3], later extended to the ABCDE criteria [4]. A lesion whose margin is smooth and regular is characteristically benign, whereas an infiltrative melanoma presents a border that is convoluted, lobulated, and uneven, reflecting the way malignant cells invade the surrounding tissue as the tumour progresses [5]. Because this cue is inherently perceptual and subject to inter-observer variation, there has been a sustained effort to replace the qualitative impression of a dermatologist with an objective, reproducible measure of border irregularity [6].

A substantial line of work has pursued this goal through boundary shape descriptors computed from dermoscopic images. The most influential of these is the fractal dimension of the lesion boundary, estimated by box-counting, which covers the contour with a sequence of grids of decreasing box size and fits the slope of the log box-count against the log inverse scale; a value close to unity indicates a smooth boundary and higher values indicate a progressively rougher, more space-filling contour. This measure has been applied to melanoma borders as a compact index of irregularity, and the prevailing expectation in the literature is that malignant lesions should exhibit a markedly elevated fractal dimension relative to benign nevi [7,8]. Alongside the fractal dimension, a family of perimeter- and hull-derived descriptors, including compactness, solidity, convexity, circularity, and asymmetry, captures complementary aspects of the same underlying shape, and automated border detection has made these measurements routine [9–12]. Machine-learning systems built on such features detect border irregularity directly [13], and deep networks operating on the whole lesion image now reach or exceed dermatologist-level accuracy on classification tasks [14–16], as systematic reviews of studies involving human experts confirm [17,18], although accuracy can fall on diverse or uncommon presentations [19]. More recent systems combine the lesion image with patient metadata [20] and extend the same machinery from diagnosis to prognosis [21]. The availability of large public reference archives such as HAM10000 and the International Skin Imaging Collaboration (ISIC) challenge collection has made these measurements reproducible at cohort scale [22–25]. What unites these approaches is their endpoint: they classify lesions or score irregularity, but they stop short of explaining which cellular process produces an irregular border in the first place.

In parallel, a mechanistic literature models tumour growth at the scale of individual cells [26–28], and recent work has begun to quantify the spatial and phenotypic heterogeneity that such models generate [29]. Hybrid cellular automata couple a discrete lattice of proliferating and migrating cells to a continuous nutrient field, and have shown that cell adhesion and contact inhibition, rather than proliferation alone, govern the invasive morphology of a growing tumour [30]. Contact inhibition is the canonical candidate for this effect: a benign cell halts division when crowded by neighbours, whereas a malignant cell that has lost this inhibition continues to divide even when surrounded, and the loss of contact inhibition is frequently invoked as the source of the finger-like, infiltrative borders seen in aggressive tumours [31,32], in which sustained proliferation and tissue invasion are recognised as core hallmarks of malignancy [33]. Off-lattice single-cell models reach a comparable conclusion in continuous space, reproducing the transition from compact monolayers to invasive aggregates as adhesion weakens [34], and direct comparisons of individual-based and continuum formulations show that the two descriptions agree on the resulting growth patterns [35]. Melanoma has itself been modelled as a multi-scale agent-based system, although such models have been directed at growth dynamics and angiogenesis rather than at the border morphology that clinicians actually observe [36]. At the boundary between the two communities, experimental observations of tumour growth interfaces have found that the rough edge of an advancing tumour obeys super-rough scaling dynamics, indicating that border roughness and growth dynamics are mechanistically coupled [37]. Together these studies establish that cell-scale rules can, in principle, determine macroscopic border shape, yet none of them closes the loop by measuring simulated and clinical borders with the same algorithm and asking which rule reproduces the observed signature.

That unclosed loop is the gap this paper addresses. The imaging literature knows that malignant borders are more irregular, but not why a specific kind of irregularity, namely lobulation, should follow from a particular cell behaviour. The modelling literature knows which rules distort a growing aggregate, but its outputs are usually assessed qualitatively, or with metrics that are not the ones a clinician would compute on a patient image. The result is a divide in which the two communities agree on the phenomenon but cannot map one onto the other, leaving the mechanistic basis of a diagnostic feature unresolved, and leaving computational pathology without a way to connect its image features to the biology that generates them, a link on which clinical translation increasingly depends [38,39].

This study closes that divide through a single quantitative pipeline applied without modification to both sides. We measured seven boundary descriptors, including the box-counting fractal dimension, on 1,581 dermoscopic lesions from the ISIC 2018 archive, and we report that the genuine malignant signature is not an elevated fractal dimension but a specific combination of shape features: higher compactness, lower solidity, and lower convexity, corresponding to a border composed of a few large lobes. We then built a hybrid cellular automaton and swept four candidate cell-level mechanisms, finding that none of them, including the widely invoked loss of contact inhibition, produces anything other than a near-circular cluster; this negative result is reported rather than tuned away. Extending the model with a long-range spatial heterogeneity in the growth rate, however, recovers the measured malignant signature to within a descriptor distance of 0.187, roughly sixfold better than the best single mechanism. A variance-based sensitivity analysis identifies the correlation length of this heterogeneity as the dominant driver by total-effect index [40–42]. The contributions of the paper are threefold. First, we establish the measured morphology of a large clinical cohort and show that the familiar fractal-dimension narrative is, on this data, the weakest of seven descriptors, whereas perimeter- and hull-derived descriptors carry the malignant signal. Second, we demonstrate, as a pre-specified negative result, that the empty-neighbour lattice assumption common to many cell models is insufficient to generate lobular borders, a finding that motivates the mechanism studied here. Third, we identify growth-field heterogeneity as the mechanism that aligns simulation to observation, and we state plainly that this field is imposed rather than emergent, thereby framing the open question of how such heterogeneity could arise from cell-autonomous dynamics. The paper moves from measurement to mechanism: we first establish the boundary signature in a large clinical cohort, then show that canonical cell-level mechanisms fail to reproduce it, and finally identify a long-range growth field that does. The methods underlying each step are described after the discussion.

## Results

### The malignant signature is lobulation, not roughness

We measured seven boundary descriptors on 1,581 lesions segmented by the automatic Otsu route and compared their median values across the three diagnostic grades (Table 1; Fig 1). The strongest and most consistent signal is a compactness that rises from a benign median of 1.351 through 1.800 in in situ melanoma to 1.810 in invasive melanoma, together with a solidity that falls from 0.910 to 0.853 and a convexity that falls from 0.936 to 0.839. A compactness well above unity and a solidity and convexity well below unity describe a border composed of a few deep indentations rather than many fine undulations, so this three-descriptor pattern is the signature of a small number of large lobes. The three descriptors separate invasive melanoma from benign nevi with area-under-the-curve values of 0.740, 0.739, and 0.738 respectively, and the between-grade difference is large and unambiguous for all three: eta-squared reaches 0.087, 0.088, and 0.086, with the benign-to-invasive pairwise comparison corrected p-values of 1.6e-12, 5.3e-12, and 4.1e-12.

**Table 1.** Median boundary descriptors by diagnostic grade on the automatic Otsu route.

| Descriptor | Benign | In situ | Invasive | $\eta^2$ | AUC | Direction |
| --- | --- | --- | --- | --- | --- | --- |
| Compactness | 1.351 | 1.800 | 1.810 | 0.087 | 0.740 | ↑ |
| Solidity | 0.910 | 0.849 | 0.853 | 0.088 | 0.739 | ↓ |
| Convexity | 0.936 | 0.841 | 0.839 | 0.086 | 0.738 | ↓ |
| Asymmetry | 0.083 | 0.131 | 0.136 | 0.075 | 0.723 | ↑ |
| Circularity | 0.863 | 0.815 | 0.826 | 0.040 | 0.664 | ↓ |
| Perimeter decay slope | -0.057 | -0.088 | -0.090 | 0.062 | 0.702 | ↓ |
| Fractal dimension | 1.040 | 1.064 | 1.048 | 0.024 | 0.625 | ≈ |

Values are medians for the automatic Otsu route ( $n = 1,581$ ). $\eta^2$ is the Kruskal–Wallis effect size and AUC is the area under the receiver-operating curve for the benign-to-invasive contrast. Direction gives the sign of the benign-to-invasive change. Compactness, solidity, and convexity carry the malignant signal; the fractal dimension is the weakest descriptor.

**Fig 1.**
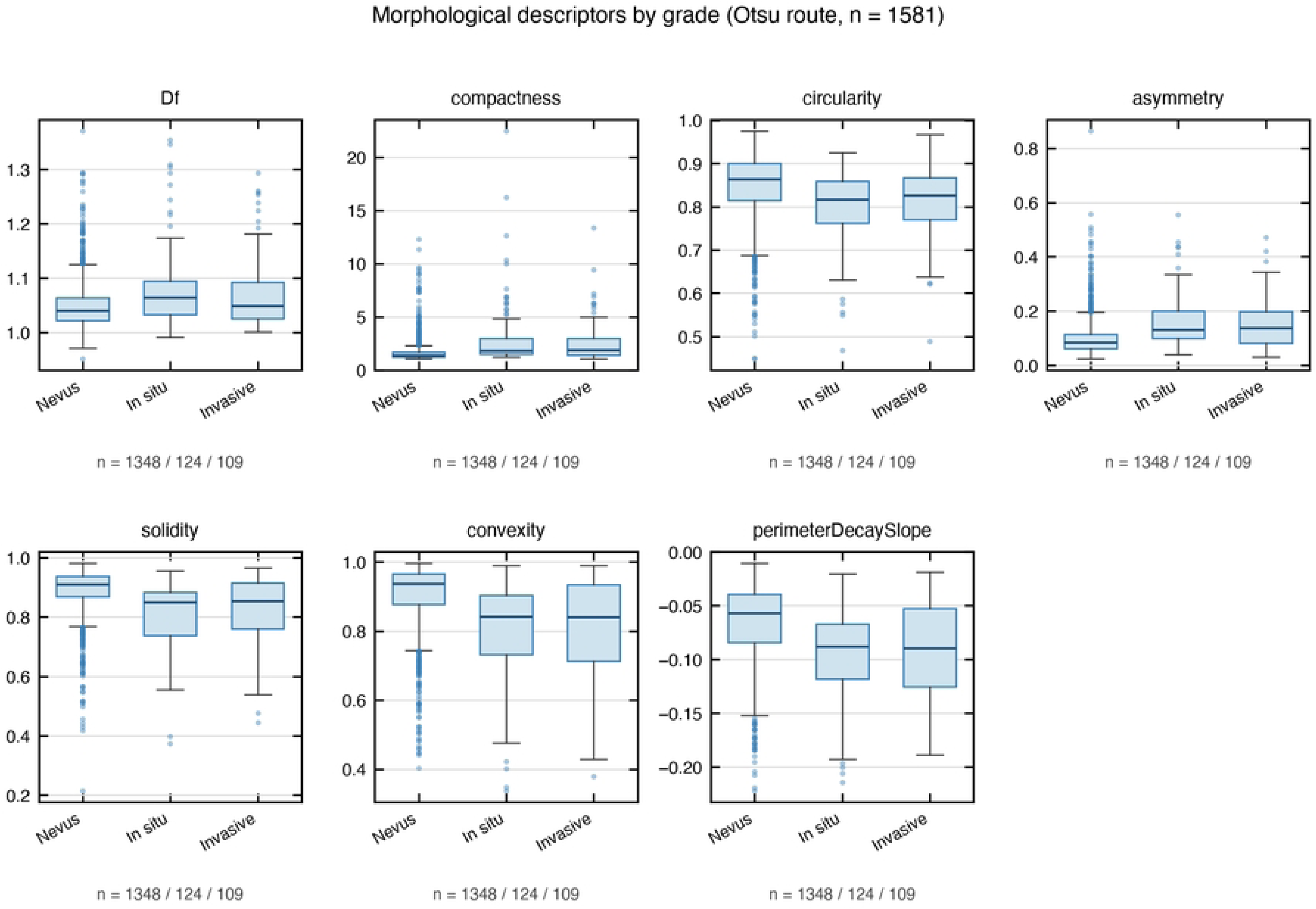
Distributions of the seven boundary descriptors by diagnostic grade. Median values are shown for the automatic Otsu route across benign, in situ, and invasive grades (n = 1,581 lesions).

The fractal dimension, by contrast, is nearly constant across grades. It moves from 1.040 in benign lesions to 1.064 in in situ melanoma and back to 1.048 in invasive melanoma, an excursion of a few hundredths, and its area under the curve of 0.625 is the weakest of the seven descriptors. Its between-grade difference is real but small, with an eta-squared of 0.024, roughly a quarter of the effect carried by any of the three primary descriptors. The fractal dimension is therefore not the carrier of the malignant signal in this cohort; the signal lives in the perimeter- and hull-derived measures rather than in the fine roughness that the fractal dimension is designed to capture. The remaining descriptors are intermediate: asymmetry rises from 0.083 to 0.136 (AUC 0.723), circularity falls from 0.863 to 0.826 (AUC 0.664), and the perimeter decay slope steepens from -0.057 to -0.090 (AUC 0.702). Asymmetry is the one intermediate descriptor that does not depend on boundary detail, being a fold overlap of the mask about its principal axis, and its effect is the largest of the three.

The direction of every effect is consistent with the lobulation account. A lesion that develops a few large lobes lengthens its perimeter relative to its area, raising compactness; it acquires indentations that reduce the filled fraction of its convex hull, lowering both solidity and convexity; and it becomes less rotationally symmetric and less circular, while its box-counting dimension barely moves because the coarse-scale lobes do not change the fine-scale slope of the box-counting line. The three grades form an ordinal severity axis, and the median trajectories of the three primary descriptors track severity: compactness climbs from benign through in situ to invasive, while solidity and convexity fall from benign to invasive, so the signature is not a simple two-class split but a graded progression that tracks severity rather than merely distinguishing the extremes of the cohort. Fig 2 shows the box-counting fits that underlie the fractal-dimension estimate, and Fig 3 shows the receiver-operating curves for all seven descriptors, where the three primary descriptors and asymmetry sit above the remaining three.

**Fig 2.**
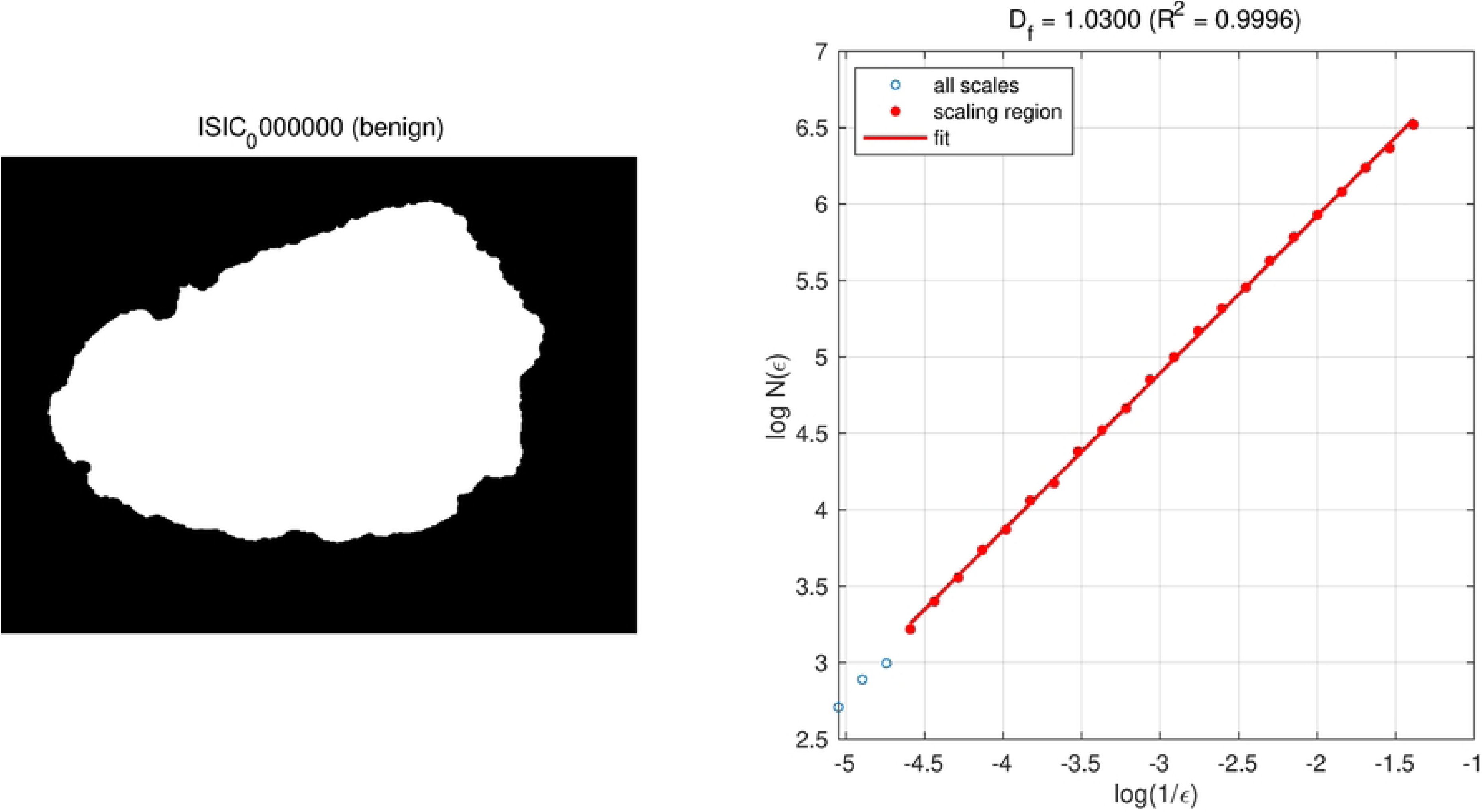
Box-counting log–log fits underlying the fractal-dimension estimate. The detected scaling region used for each fit is indicated.

**Fig 3.**
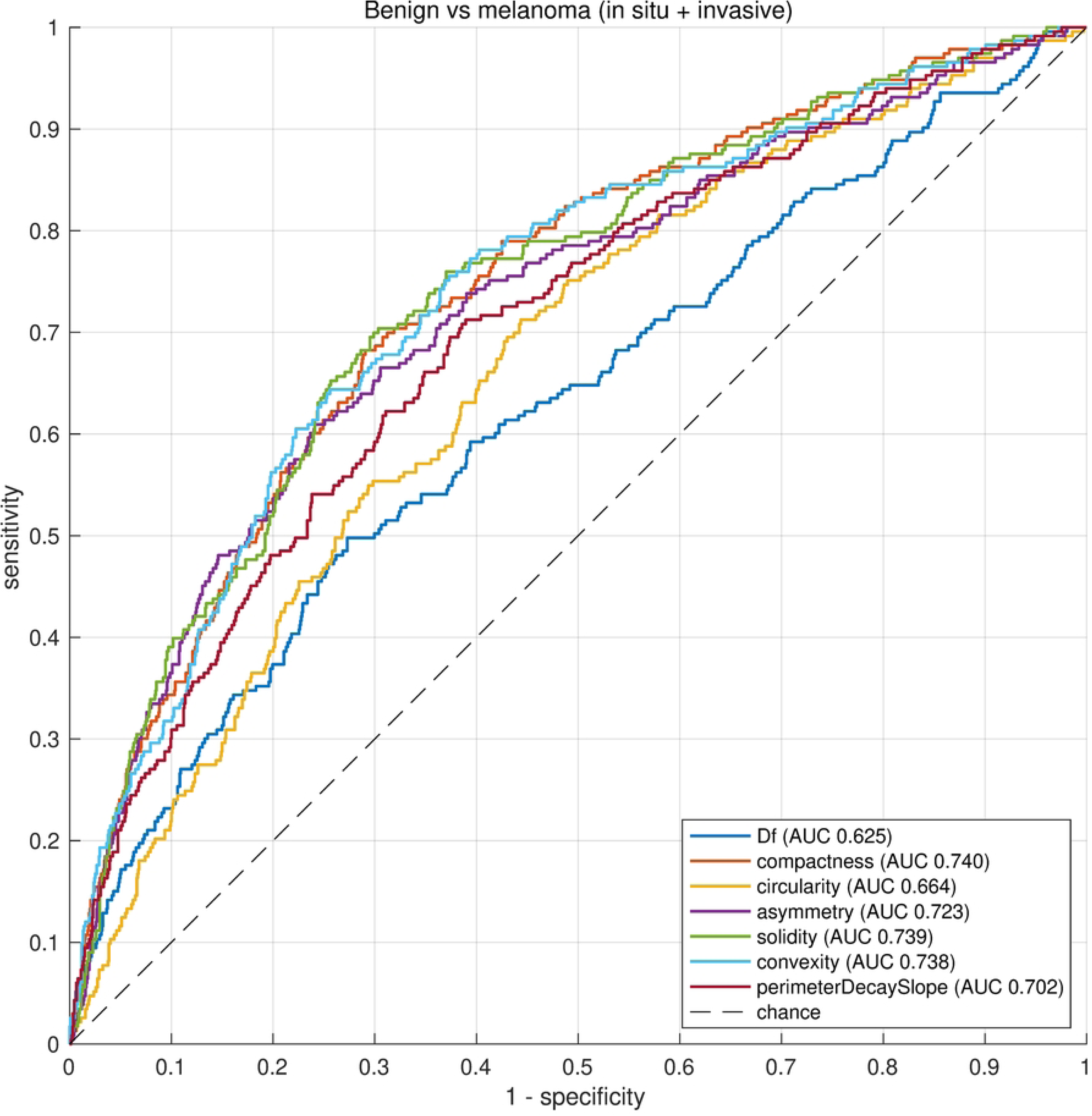
Receiver-operating curves for the seven descriptors separating invasive melanoma from benign nevi. Area-under-the-curve values are reported in Table 1.

### The signature survives the segmentation-route test

A between-grade difference in shape descriptors could in principle be a difference in segmentation quality rather than in shape, so we repeated the entire measurement on the expert ground-truth route and on two specific confounds. Each route was analysed on the lesions it accepted, 2,088 for the expert masks and 1,581 for the automatic masks. The result is a documented failure of the expert route rather than a confirmation of it. On the expert route, five of the seven descriptors run in the direction opposite to biological expectation: invasive melanoma measures less compact, more solid, and more convex than benign nevi, and the perimeter decay slope and fractal dimension also reverse (Table 2). All five reversals survive Holm correction, with adjusted p-values from 8.6e-05 to 1.4e-02, but with small effect sizes, a rank-biserial of 0.14 to 0.21 and an eta-squared of 0.003 to 0.009. The honest reading is that the bias is systematic and reproducible, not that it is large.

**Table 2.** Segmentation-route test of descriptor direction on the expert and automatic routes.

| Descriptor | Expected direction | Expert $r_{rb}$ | Expert $p_{adj}$ | Otsu $r_{rb}$ | Otsu $p_{adj}$ |
| --- | --- | --- | --- | --- | --- |
| Compactness | + | -0.156 | $5.5 \times 10^{-3}$ | 0.408 | $1.6 \times 10^{-12}$ |
| Convexity | - | 0.192 | $3.5 \times 10^{-4}$ | -0.400 | $4.1 \times 10^{-12}$ |
| Solidity | - | 0.142 | $1.4 \times 10^{-2}$ | -0.399 | $5.3 \times 10^{-12}$ |
| Perimeter decay slope | - | 0.208 | $8.6 \times 10^{-5}$ | -0.342 | $4.7 \times 10^{-9}$ |
| Fractal dimension | + | -0.186 | $5.5 \times 10^{-4}$ | 0.192 | $1.7 \times 10^{-3}$ |
| Asymmetry | + | 0.133 | $1.7 \times 10^{-2}$ | 0.384 | $3.2 \times 10^{-11}$ |
| Circularity | - | -0.072 | 0.22 (n.s.) | -0.293 | $5.8 \times 10^{-7}$ |

$r_{rb}$ is the rank-biserial correlation and $p_{adj}$ the Holm-adjusted p-value. Five of seven descriptors reverse on the expert route; all seven hold their expected direction on the automatic route. n.s., not significant.

The mechanism of the reversal is annotation coarseness, not missing detail. The ISIC 2018 ground-truth masks are substantially polygon-traced: the median fraction of boundary vertices lying on a straight run is 0.587, and 24.7 percent of masks exceed 0.85, that is, they are effectively hand-drawn polygons. A polygon boundary is locally straight by construction, so it attenuates any boundary-detail descriptor; but attenuation can only push an effect toward zero, never past it. A sign reversal therefore requires an active mechanism, and the most plausible one is that annotation coarseness scales with lesion size, so that larger, more irregular lesions receive proportionally coarser polygons and end up smoother than small benign nevi. The data agree with that prediction: the two descriptors that do not reverse are exactly the two that do not depend on boundary detail, namely asymmetry and circularity, while all five reversed descriptors depend on perimeter length, concavity, or the fine boundary itself. On the automatic route, by contrast, all seven descriptors point in the expected direction, all seven are significant, and the effect sizes are roughly an order of magnitude larger for the perimeter- and hull-derived measures.

The contrast between the two routes is sharpest in effect size: on the automatic route the eta-squared values for the perimeter- and hull-derived descriptors span 0.024 to 0.088 with area-under-the-curve values of 0.62 to 0.74, whereas on the expert route the same descriptors span 0.003 to 0.009 with area-under-the-curve values of 0.52 to 0.61. The automatic route therefore does not merely preserve the sign of the effect; it carries roughly an order of magnitude more of it. We then tested the automatic route’s advantage against the confound that matters, a segmentation-quality gradient that co-varies with severity. Agreement between the two routes does decline with severity, with a Jonckheere–Terpstra z of -3.38 and a p of 7.3e-04, so the concern is real and must be met. Restricted to well-segmented lesions only, defined as a Dice overlap of at least 0.75 and numbering 1,232 paired lesions, all seven descriptors remain significant with p-values from 5.4e-06 to 9.8e-22 and only modest attenuation of the effect sizes. Under the artefact explanation the effects would collapse in this subset; they do not. A residual-hair confound was tested in the same spirit: the fractal dimension does rise with hair fraction, with a Spearman correlation of 0.168, and hair burden differs by grade, yet restricted to the least-hairy forty percent the fractal dimension’s effect size nearly doubles rather than vanishes, so hair masks the signal rather than manufacturing it. Figs 4–7 present the Bland–Altman agreement, the by-grade Dice agreement, the perimeter scale stability, and the sign-reversal divergence respectively.

**Fig 4.**
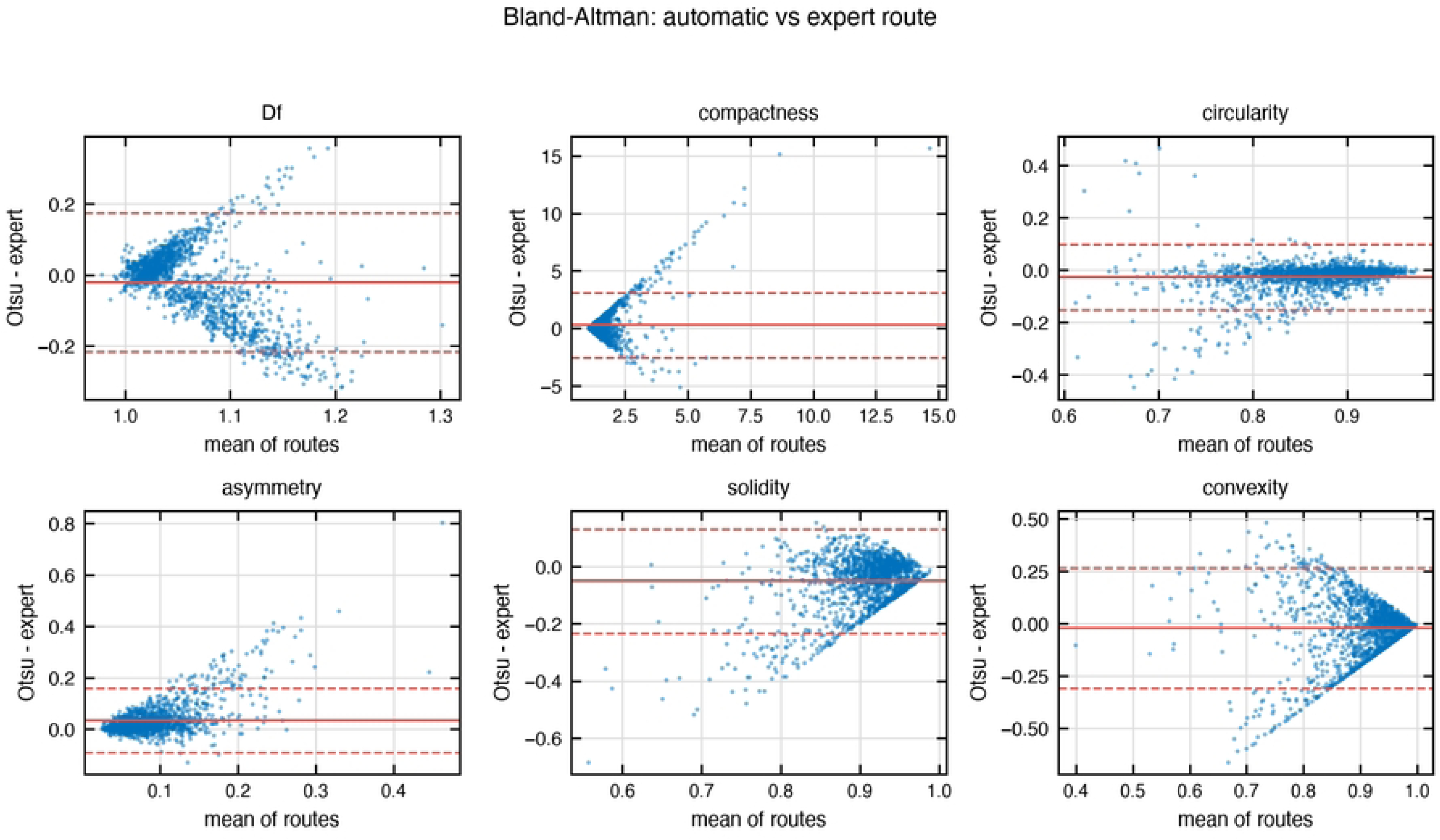
Bland–Altman agreement between the expert and automatic segmentation routes. Differences are shown against the mean of the two routes.

**Fig 5.**
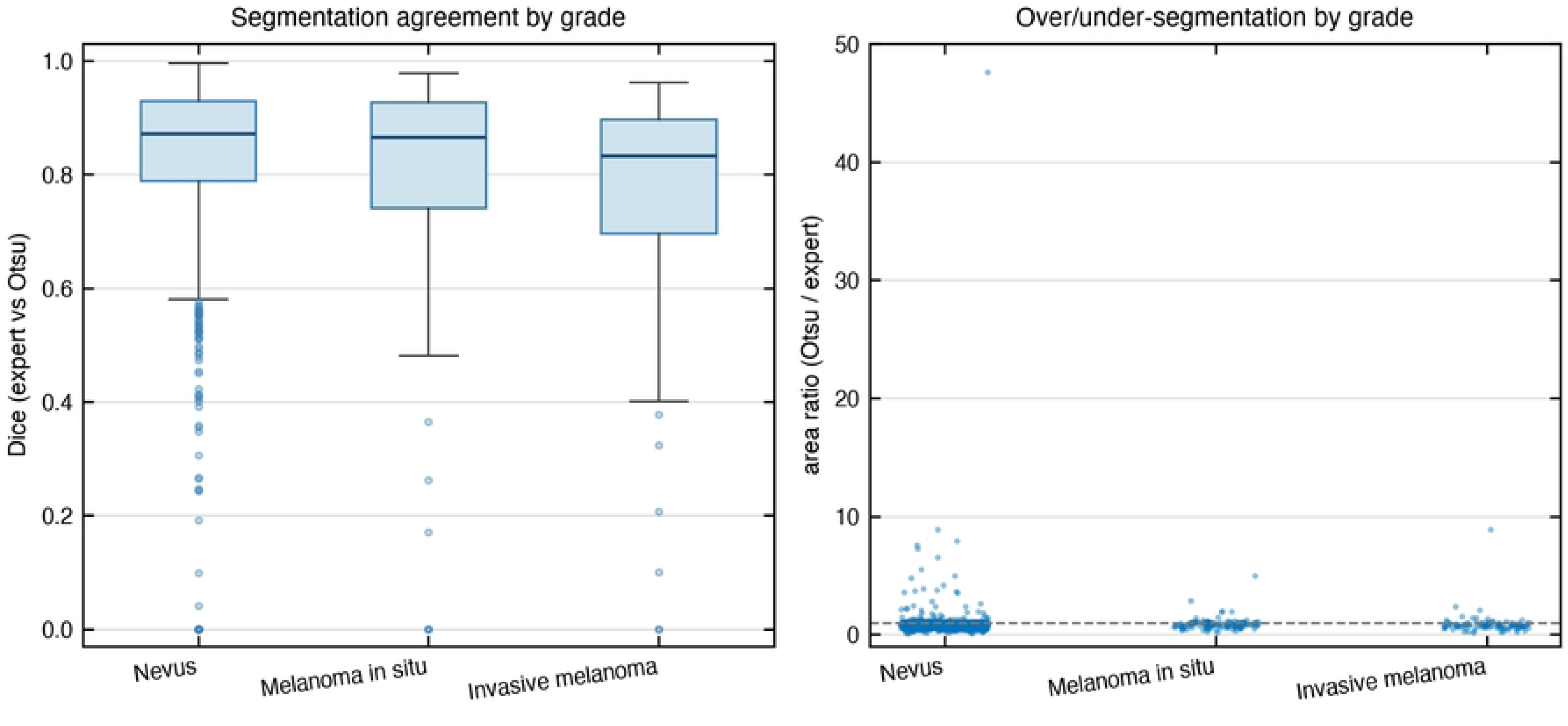
Dice agreement between the two segmentation routes by diagnostic grade. Agreement declines with severity, a trend tested with the Jonckheere–Terpstra test.

**Fig 6.**
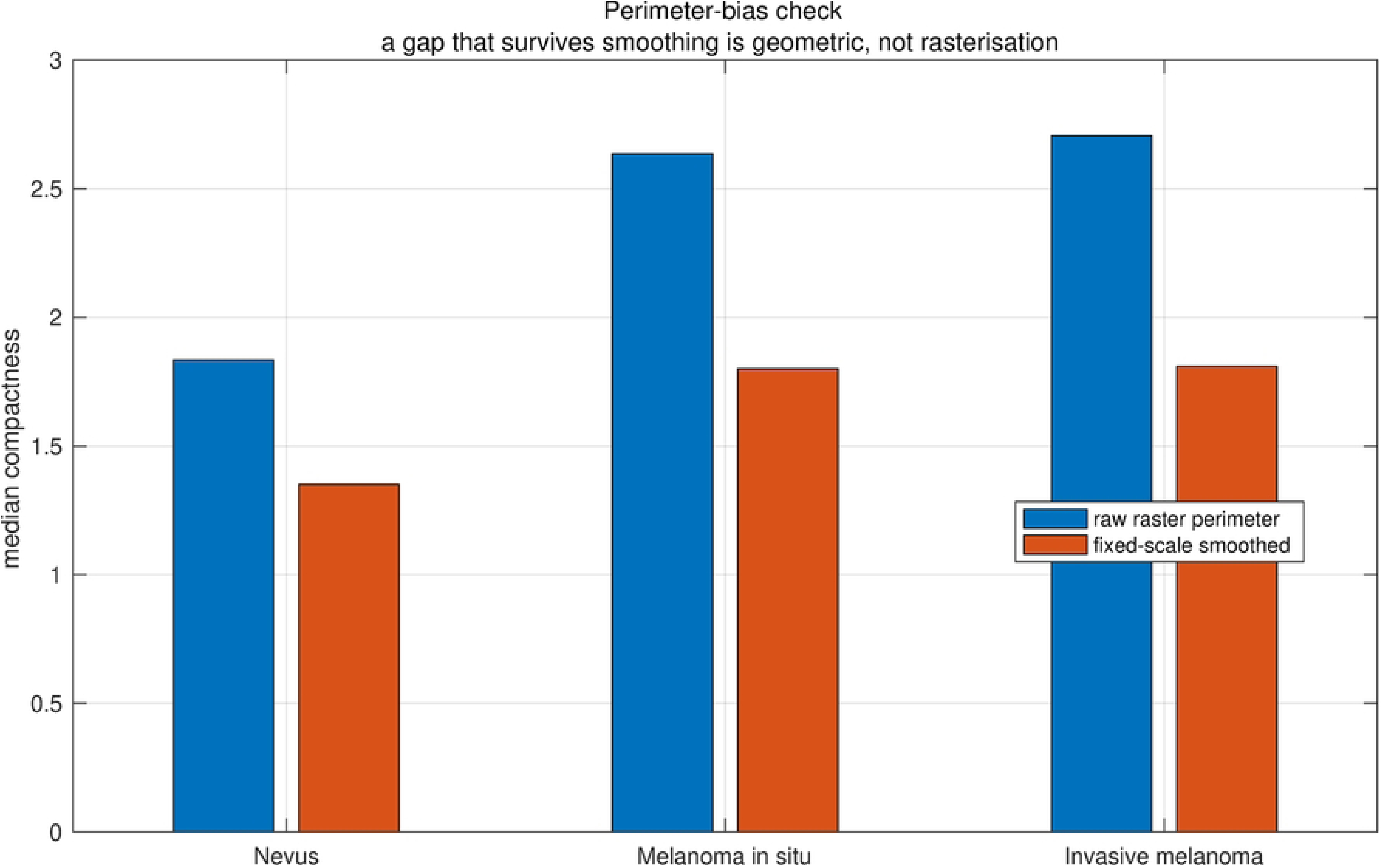
Perimeter scale stability of the smoothed lesion boundary. Perimeter is plotted against the smoothing scale for the automatic route.

**Fig 7.**
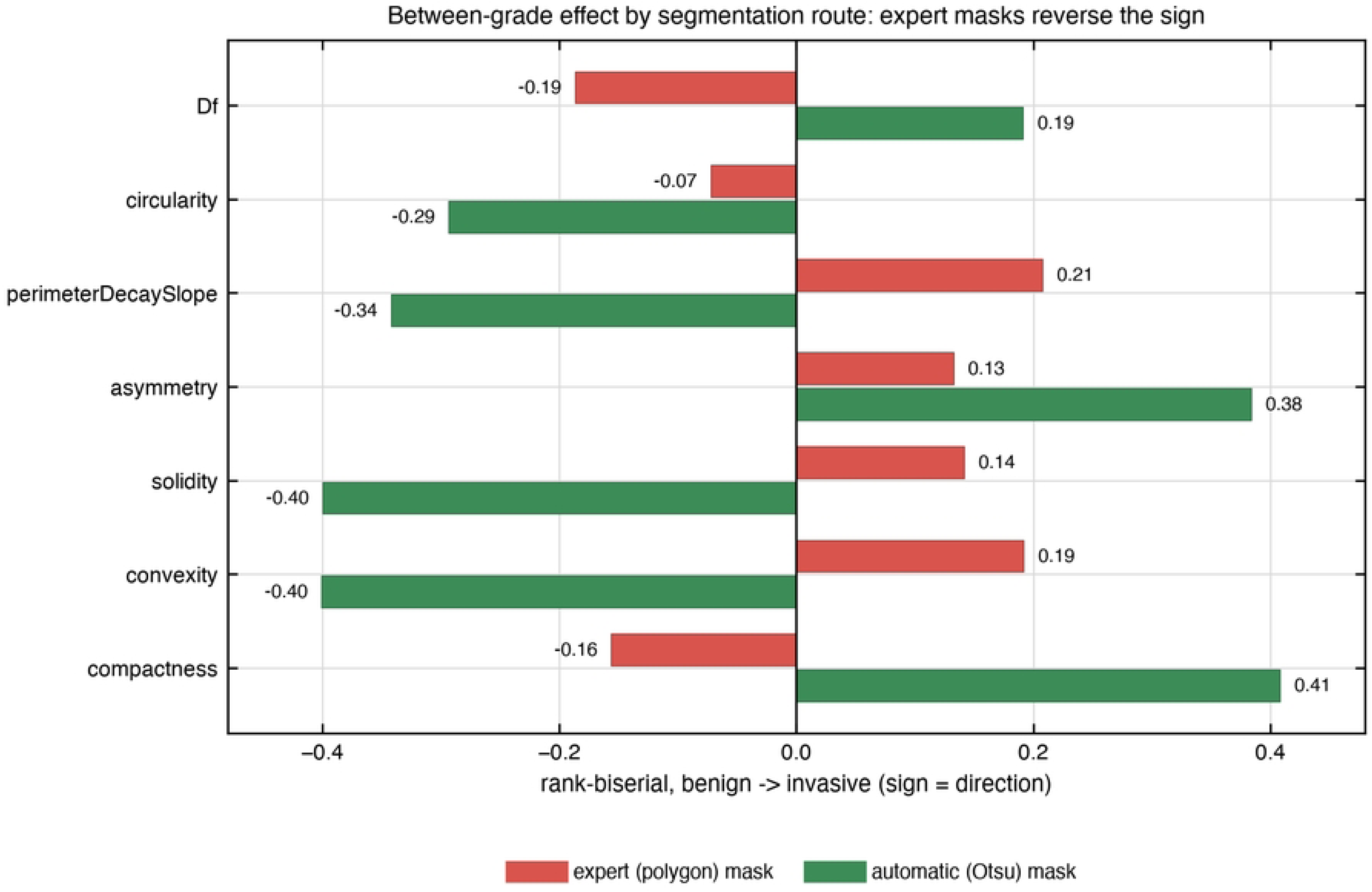
Divergence of descriptor sign between the expert and automatic routes. Five of seven descriptors reverse on the expert route.

### Four canonical mechanisms fail to escape the circle

We next asked which cell-level rule could produce the measured lobular signature, sweeping four canonical mechanisms in the hybrid cellular automaton, each at five levels with three seeds. The answer is none of them. At their best parameter settings, all four produce near-circular clusters whose distance from the median invasive-melanoma target in the space of compactness, solidity, and convexity lies between 1.147 and 1.166 (Table 3; Fig 8), and the winning setting for each arm sits at a mild or mid-range level rather than at an extreme, so the failure is not for want of an aggressive sweep. Loss of contact inhibition and clonal heterogeneity both stall at a distance of 1.166, enhanced migration at 1.153, and nutrient fingering, the closest, at 1.147. The simulated boundaries all sit at a compactness of roughly 1.02 to 1.03, a solidity near 0.98, and a convexity near 0.99, whereas the malignant target sits at 1.810, 0.853, and 0.839. Concretely, the best clonal-heterogeneity and contact-inhibition runs reach a compactness of 1.023, a solidity of 0.977, and a convexity of 0.994; the best enhanced-migration run reaches 1.030, 0.976, and 0.992; and the best nutrient-fingering run reaches 1.035, 0.975, and 0.992. In other words, no combination of these mechanisms moves the simulated tumour more than a few percent away from a perfect circle.

**Table 3.** Best configuration of each canonical mechanism and its distance to the malignant target.

| Mechanism | Level | Seed | Distance | Compactness | Solidity | Convexity |
| --- | --- | --- | --- | --- | --- | --- |
| Loss of contact inhibition | 1 | 3 | 1.166 | 1.023 | 0.977 | 0.994 |
| Clonal heterogeneity | 1 | 3 | 1.166 | 1.023 | 0.977 | 0.994 |
| Enhanced migration | 3 | 1 | 1.153 | 1.030 | 0.976 | 0.992 |
| Nutrient fingering | 2 | 2 | 1.147 | 1.035 | 0.975 | 0.992 |

Distance is measured in the z-scored space of compactness, solidity, and convexity from the median invasive-melanoma target (compactness 1.810, solidity 0.853, convexity 0.839). All four mechanisms remain within a few percent of a perfect circle.

**Fig 8.**
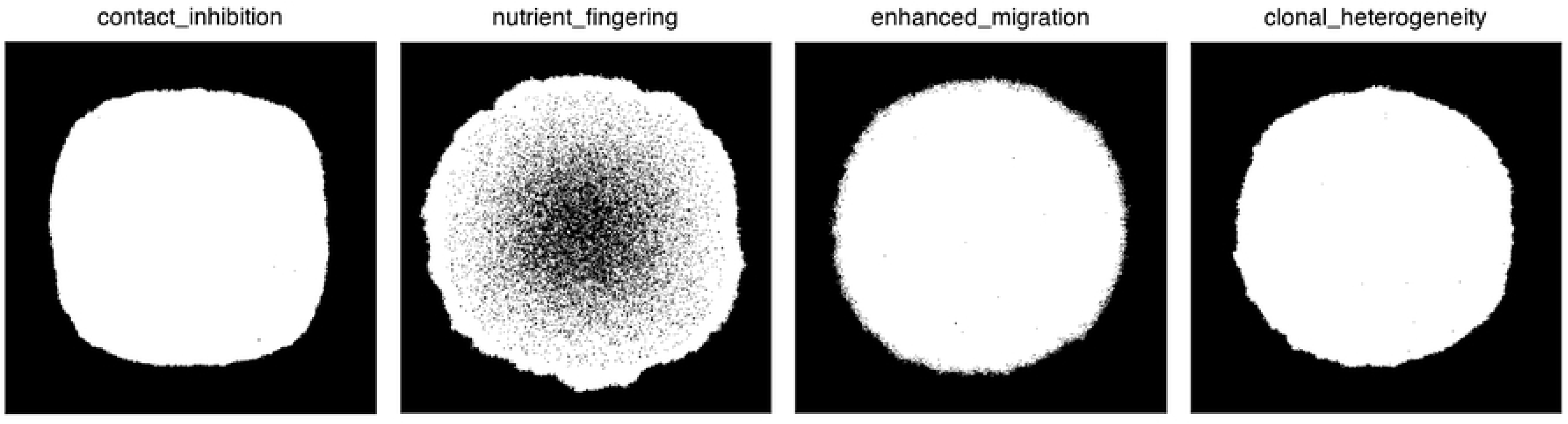
Representative morphologies produced by the four canonical mechanisms. All four arms produce near-circular clusters rather than lobulated borders.

The reason is structural rather than a failure of tuning. In every arm, a cell can only divide into an empty neighbouring site, so a cell surrounded by occupied neighbours has nowhere to place a daughter and becomes quiescent. Losing contact inhibition, for example, only changes how strongly crowding blocks division; it does not create the empty sites that division requires, so the tumour simply smooths its own frontier faster. Clonal heterogeneity, enhanced migration, and nutrient fingering likewise modify the rate or direction of growth without breaking the empty-neighbour bottleneck, and all four therefore converge on the same Eden-like near-circular cluster. The negative result is informative precisely because it is robust across four independent mechanisms: the empty-neighbour lattice assumption common to this class of model is itself insufficient to generate lobular borders. Fig 9 shows the trajectories of the four arms through descriptor space, each of which remains confined to the near-circular corner far from the malignant target. This result was not tuned away; it is reported as the motivation for the mechanism tested next.

**Fig 9.**
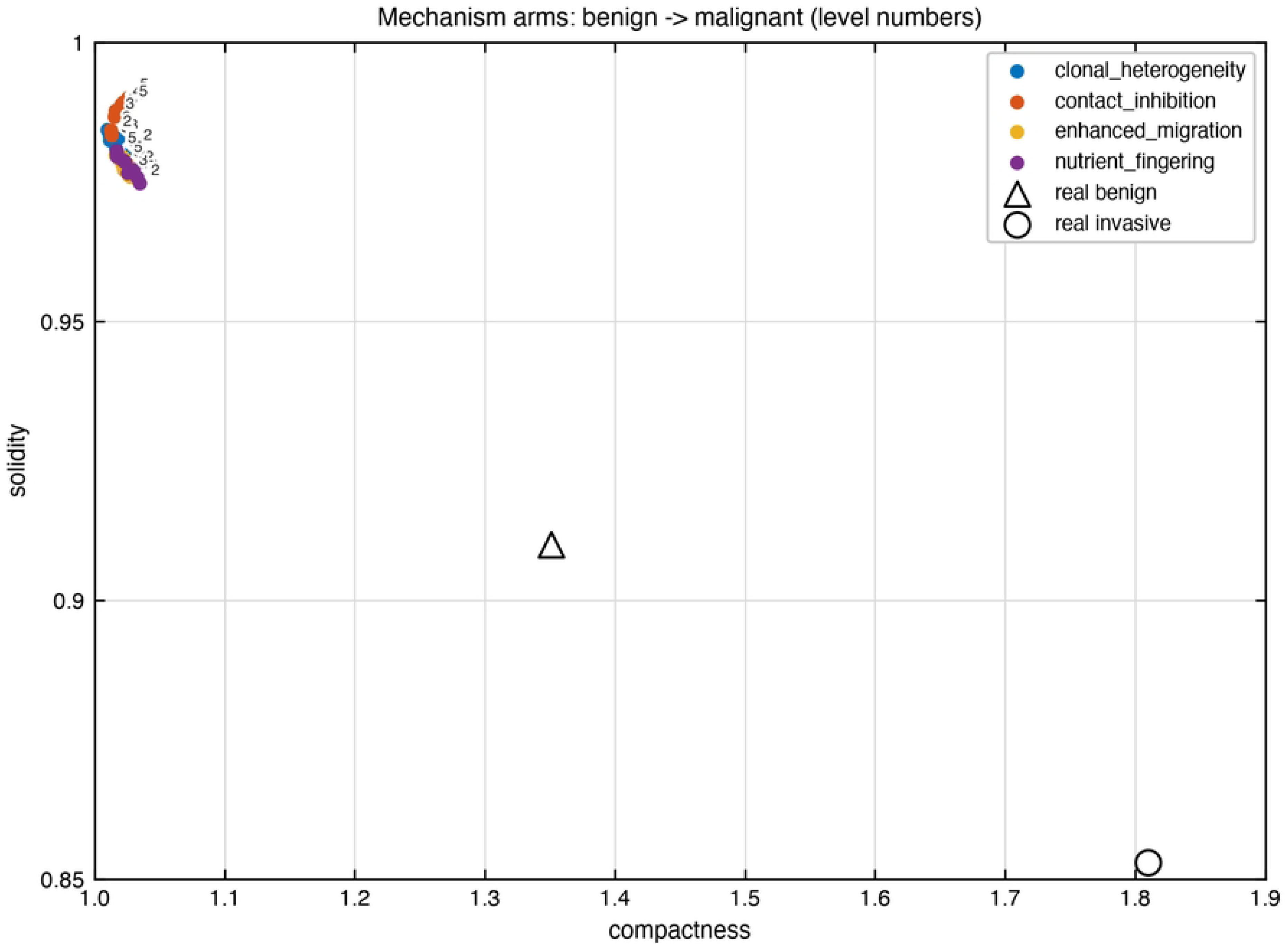
Trajectories of the four canonical mechanisms through descriptor space. Each arm remains confined to the near-circular corner, far from the malignant target.

### A long-range growth field aligns simulation to observation

Because the four canonical mechanisms failed on the same assumption, the alignment analysis lifted that assumption in two different ways and compared them directly. The first, mechanical displacement, lets a surrounded cell push a neighbouring cell into an empty site, so division no longer strictly requires an adjacent vacancy. The second, spatial heterogeneity, multiplies each cell’s division rate by a smooth static growth-rate field that varies over a long correlation length, so different regions of the same tumour proliferate at different rates. In the coarse sweep, the two mechanisms were compared over five levels with three seeds (Fig 10). Mechanical displacement converges back to a near-circular cluster, with a best distance of 1.125, no better than the four canonical mechanisms. Spatial heterogeneity is different: its best coarse-level distance is 0.406, already a threefold improvement, and it produces a compactness of 1.485, a solidity of 0.886, and a convexity of 0.902, the first simulated boundary to move substantially off the circle.

**Fig 10.**
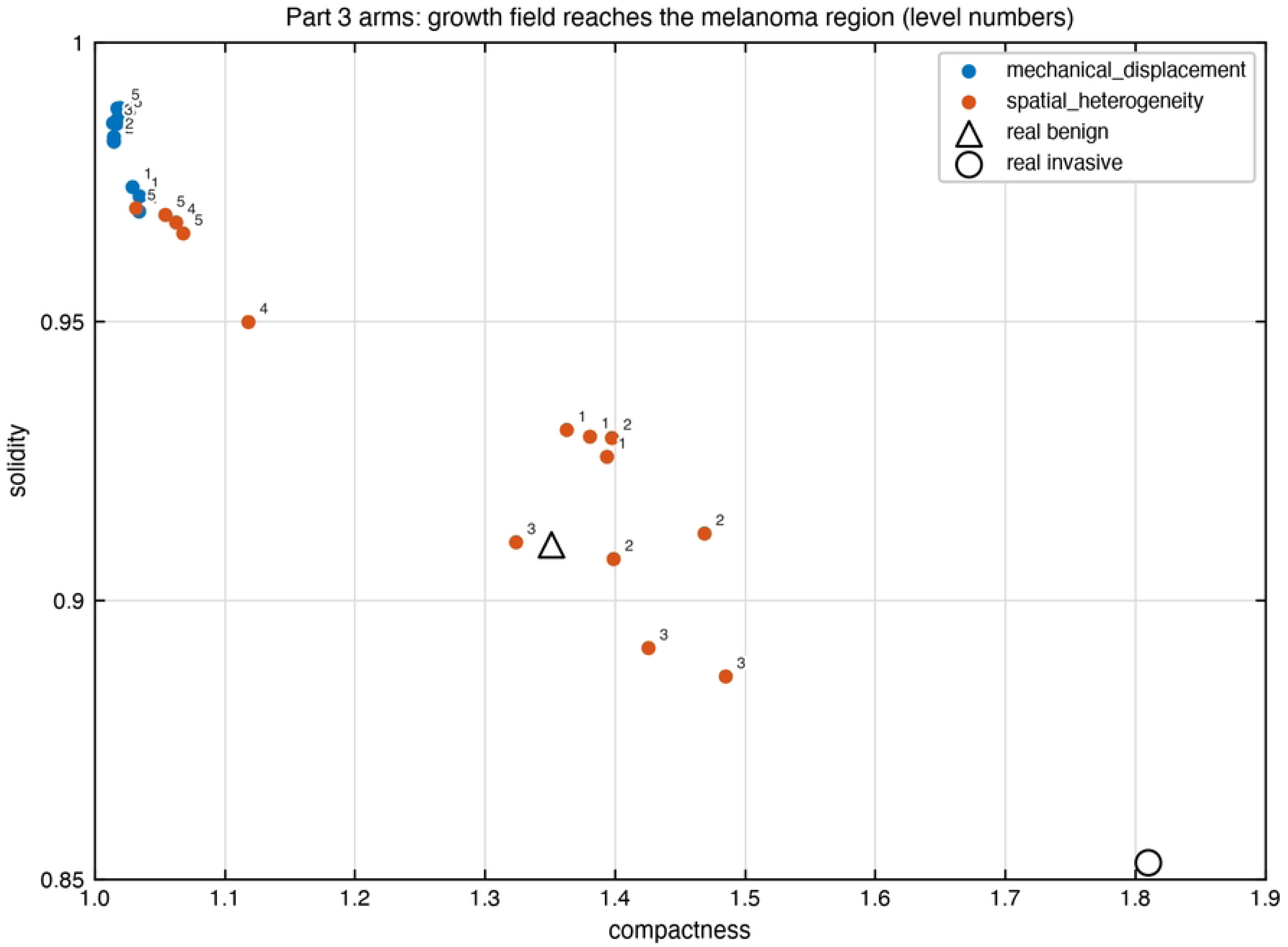
Coarse-sweep trajectories of mechanical displacement and spatial heterogeneity. Spatial heterogeneity moves substantially toward the malignant target; mechanical displacement does not.

The refinement stage combined a one-at-a-time screening, a variance-based sensitivity analysis, and a two-dimensional phase diagram over the field amplitude and correlation length (Fig 11; Table 4). The optimum lies at an amplitude of 0.58 and a correlation length of 58 pixels, and there the three-descriptor distance falls to 0.187. The best simulated boundary has a compactness of 1.835 against a target of 1.810, a solidity of 0.873 against 0.853, and a convexity of 0.809 against 0.839, so the simulated lesion reproduces the measured malignant signature to within a few percent on all three primary descriptors simultaneously. The alignment is roughly sixfold better than the best of the four canonical mechanisms. The sensitivity analysis (Fig 12) shows that the correlation length dominates: its first-order index is 0.113 and its total-effect index is 0.892, while the amplitude carries a first-order index of 0.257 and a total-effect index of 0.406, from 1,536 evaluations with a base sample of 256 points over the swept ranges of 0.1 to 1.0 in amplitude and 24 to 284 pixels in correlation length. The large gap between the correlation length’s first-order and total-effect indices reflects its interaction with the amplitude, meaning the two parameters must be set jointly to reach the optimum. The phase diagram confirms the structure: the low-distance region is not a single point but a band in the parameter plane, and the alignment distance rises steeply when the correlation length is either too short, collapsing back toward the near-circular cluster, or too long, where the field ceases to modulate the boundary at the lesion scale. The one-at-a-time screening shows the two parameters to be of comparable individual influence, with a maximum absolute change in the alignment distance of about 0.31 over each swept range, which is consistent with the sensitivity analysis’s indication that the parameters act jointly rather than additively. The winning mechanism is an imposed field, not an emergent instability, and the model states this explicitly; the open question of how such a long-range growth-rate heterogeneity could arise from cell-autonomous dynamics is taken up in the discussion.

**Fig 11.**
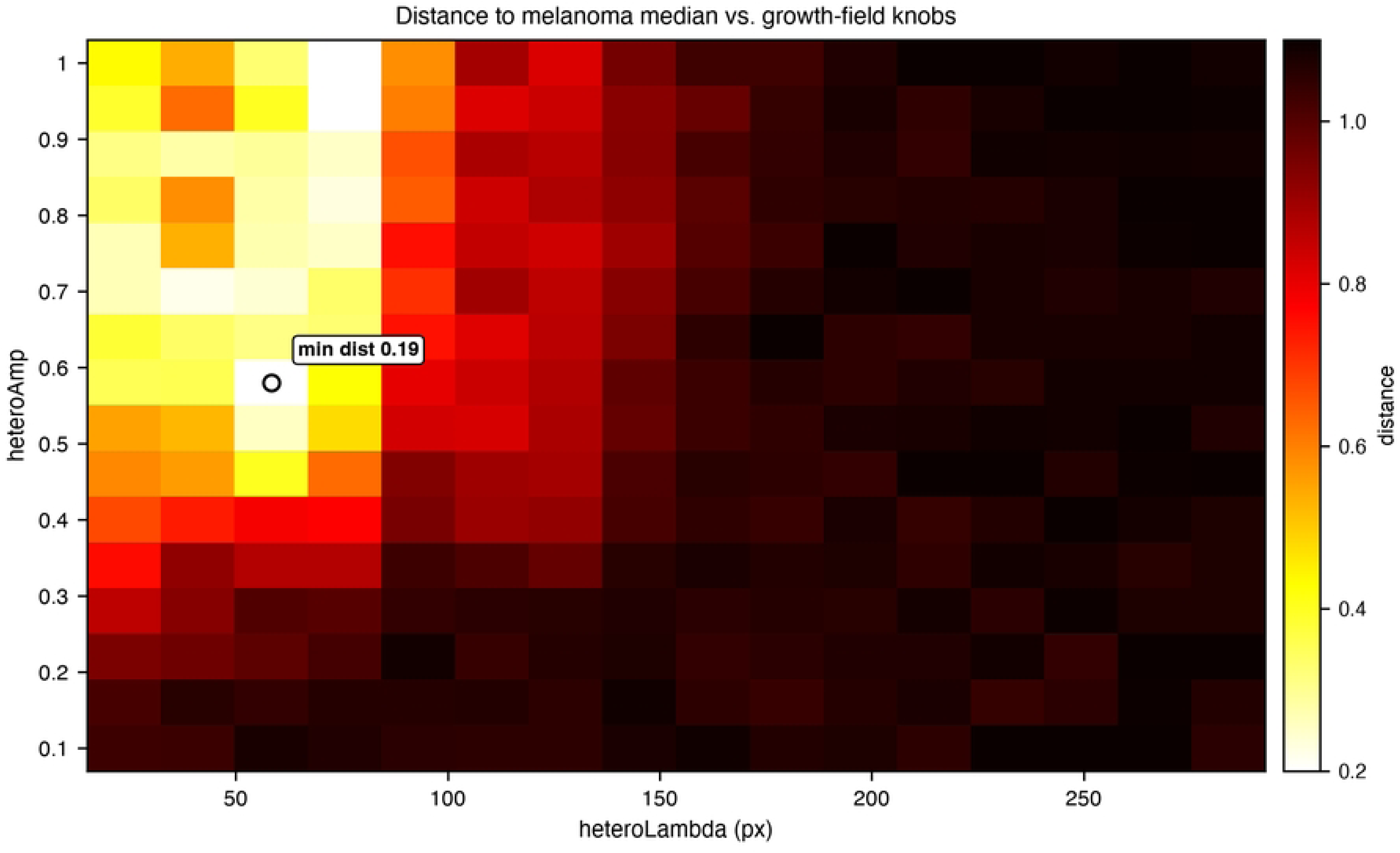
Phase diagram of alignment distance over field amplitude and correlation length. The low-distance region forms a band rather than a single optimum point.

**Table 4.** Alignment outcomes and variance-based sensitivity indices for the growth-rate field.

| Quantity | Value |
| --- | --- |
| Mechanical displacement, coarse sweep, best distance | 1.125 |
| Spatial heterogeneity, coarse sweep, best distance | 0.406 |
| Optimum amplitude | 0.58 |
| Optimum correlation length | 58 px |
| Optimum three-descriptor distance | 0.187 |
| Optimum compactness | 1.835 |
| Optimum solidity | 0.873 |
| Optimum convexity | 0.809 |
| Amplitude, first-order Sobol' index ($S_i$) | 0.257 |
| Amplitude, total-effect Sobol' index ($S_{Ti}$) | 0.406 |
| Correlation length, first-order Sobol' index ($S_i$) | 0.113 |
| Correlation length, total-effect Sobol' index ($S_{Ti}$) | 0.892 |

Distances are measured in the z-scored space of compactness, solidity, and convexity. The malignant target is compactness 1.810, solidity 0.853, and convexity 0.839. Sobol' indices come from 1,536 evaluations with a base sample of 256 points over amplitude 0.1–1.0 and correlation length 24–284 px.

**Fig 12.**
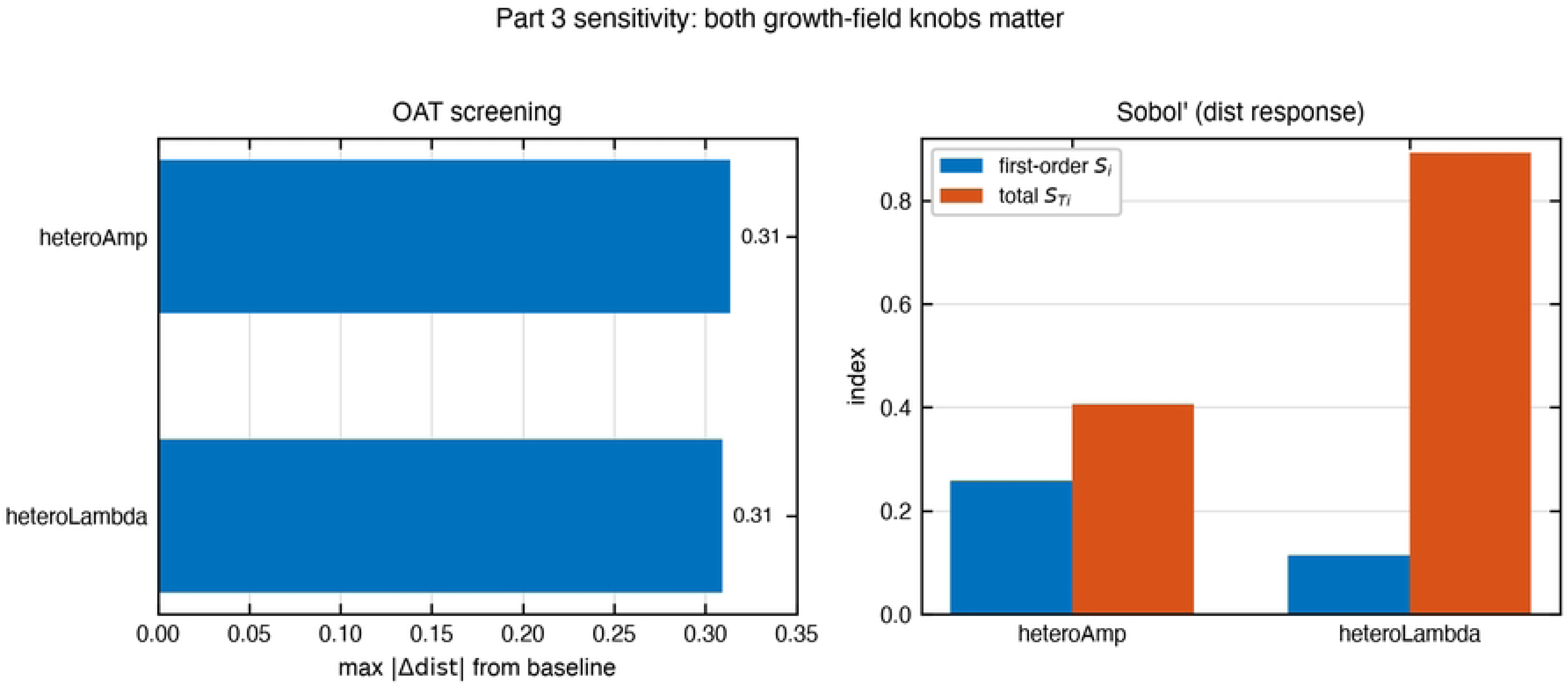
Variance-based sensitivity indices of the growth-rate field parameters. Correlation length carries the larger total-effect index, indicating spatial structure rather than growth rate as the decisive property.

## Discussion

### A correction to the fractal-dimension narrative

The first result of this study is a correction to a widely held expectation. A substantial line of work has used the box-counting fractal dimension of the lesion boundary as the canonical index of border irregularity, and the prevailing expectation is that malignant lesions should show a markedly elevated fractal dimension relative to benign nevi [6,7]. On a cohort of 1,581 automatically segmented lesions we find the opposite emphasis: the fractal dimension is the weakest of seven descriptors, moving only between 1.040 in benign lesions, 1.064 in situ, and 1.048 in invasive melanoma, and carrying an area under the curve of 0.625, while the malignant signal is carried by compactness, solidity, and convexity, whose area-under-the-curve values are 0.740, 0.739, and 0.738. The signature these three descriptors jointly describe is not fine roughness but a border composed of a few large lobes. This is not a refutation of the fractal-dimension method so much as a redirection of it: the distinction between a smooth and an irregular margin is real, but the irregularity that separates melanoma from nevus in this cohort is coarse-scale lobulation, which is invisible to the fine-scale slope that box-counting estimates. The two findings are compatible once scale is made explicit, and they jointly suggest that the literature’s focus on the fractal dimension has under-weighted the perimeter- and hull-derived measures that actually carry the diagnostic signal [13].

### The segmentation-route lesson

The second result is methodological and generalises beyond melanoma. When the identical measurement pipeline is run on the polygon-traced expert masks of the same archive, five of seven descriptors reverse sign, and the two that survive are exactly the two that do not depend on boundary detail. Because attenuation can only shrink an effect toward zero and never push it past zero, the reversal is not a failure of detail but an active artefact of annotation coarseness that scales with lesion size. The practical consequence is that any study that derives perimeter- or hull-based shape features from hand-traced boundaries is at risk of measuring its own annotation protocol rather than the lesion, and the sign of that risk is correlated with severity, which is precisely the direction a diagnostic study least wants. We therefore treat the automatic Otsu route as primary throughout, not because it is perfect but because its errors are closer to uniform across grades and are testable; the expert route is retained as a sensitivity arm whose documented failure is itself evidence. This is a concrete argument for the reproducibility-motivated choices that the automatic route encodes, such as computing the Otsu threshold inside the dermatoscope aperture and leaving morphological opening disabled, and it is a warning for the wider community that ground-truth masks are not automatically ground truth for boundary geometry.

### The empty-neighbour assumption is the bottleneck

The third result is a pre-specified negative result with a positive interpretation. Four canonical cell-level mechanisms, loss of contact inhibition, clonal heterogeneity, enhanced migration, and nutrient fingering, were each swept over five levels and three seeds, and every one of them converged on a near-circular cluster, the best of them at a three-descriptor distance of 1.147 from the malignant target. The failure is structural rather than a failure of tuning. In every arm a cell can divide only into an empty neighbouring site, so a cell that is already surrounded has nowhere to place a daughter and becomes quiescent; weakening contact inhibition changes how strongly crowding blocks division, but it does not create the empty site that division requires. This result bears directly on a modelling literature that has long invoked loss of contact inhibition as the source of infiltrative, finger-like borders in aggressive tumours [30]. Our data do not show that contact inhibition is irrelevant; they show that, inside a model whose division rule assumes an empty neighbour, contact inhibition alone cannot produce the observed lobulation because the bottleneck lies one step earlier, in the assumption itself. The negative result is thus diagnostic: it localises the missing ingredient to the empty-neighbour lattice rule that this class of hybrid cellular automaton inherits, and it motivates the mechanism we tested next rather than a wider parameter sweep that would have failed for the same reason.

### A long-range growth field aligns simulation to observation

The fourth result closes the loop. Lifting the empty-neighbour assumption in two different ways, we find that mechanical displacement, which lets a surrounded cell push its neighbour aside, still converges to a near-circular cluster at a distance of 1.125, whereas a smooth, long-range spatial heterogeneity in the growth rate recovers the measured malignant signature to a three-descriptor distance of 0.187. At the optimum the simulated boundary reaches a compactness of 1.835 against an observed 1.810, a solidity of 0.873 against 0.853, and a convexity of 0.809 against 0.839, reproducing all three primary descriptors simultaneously to within a few percent. The sensitivity analysis is decisive about which knob matters: the correlation length of the growth-rate field carries a total-effect index of 0.892 while its amplitude carries 0.406, so the mechanism is one of spatial structure rather than of raw growth rate. This is the central finding of the study, and it is stated with full awareness of its limitation. The winning field is imposed on the model, not generated by it; the model demonstrates that a growth-rate field with a correlation length on the order of a hundred micrometres to millimetres is sufficient to produce lobular borders, but it does not yet explain how such a field would arise from cell-autonomous dynamics. That distinction between imposed and emergent is the open question the study hands forward, and it is deliberately not elided by relabelling the imposed field as an emergent instability.

### Relation to the tumour-growth literature

The finding connects to an older observation about the rough edge of advancing tumours, which has been reported to obey super-rough scaling dynamics and to couple boundary roughness to growth dynamics [37]. Our results are consistent with a coupling between growth and boundary shape, but they assign the coupling to a coarse, long-range heterogeneity rather than to the fine-scale roughening that fractal analysis would capture. They are also consistent with off-lattice single-cell models that tie invasive morphology to weakened adhesion, in the sense that both our mechanical-displacement arm and those models show that local mechanical rules alone tend toward compact or mildly irregular aggregates [34], and with comparisons showing that individual-based and continuum descriptions of the same process agree on the resulting morphology [35]; what moves the simulated boundary off the circle in our study is a spatially structured, non-local growth signal rather than a local interaction. Melanoma has previously been modelled as a multi-scale agent-based system directed at growth dynamics and angiogenesis [36]; our contribution is to point that same modelling machinery at the border morphology a clinician actually observes and to score its output with the same descriptors computed on patient images, which is the step that makes the mapping between image feature and biology quantitative rather than qualitative.

## Limitations and the path to emergence

Several limitations bound the conclusions. The model is two-dimensional and treats the lesion as a growing aggregate rather than a three-dimensional tumour with a dermis beneath it, so the correlation length reported here should be read as an effective two-dimensional quantity rather than a physical measurement. The cohort is a single public archive, ISIC 2018, with its own acquisition and annotation idiosyncrasies, and although the dual-route analysis argues that the signature is not a segmentation artefact of that archive, an independent cohort would strengthen the claim of generality. The grade imbalance, with 1,348 benign lesions against 109 invasive, is inherent to the source and was handled with non-parametric statistics and bootstrap intervals, but it limits the precision of the invasive-grade estimates. The fractal dimension is a single estimator, box-counting, and while its scaling-region detection was designed to avoid end truncation, other estimators such as the capacity or correlation dimension might behave differently. Finally, the decisive mechanism is imposed rather than emergent, and this is the most important limitation of all, because it is also the clearest direction for future work.

The path from imposed to emergent is the natural next step, and the study was designed to make that step concrete. A growth-rate field with a long correlation length could plausibly arise from heritable differences among subclones in the carrying capacity of their local environment [43,44], from a metabolic or nutrient gradient imposed by the surrounding tissue, or from a reaction-diffusion feedback of the kind Turing described, in which proliferation and a secreted factor jointly generate spatial structure [45]. The sensitivity result already tells us what such a mechanism must achieve: it must produce spatial structure whose dominant scale is the correlation length, not the growth rate itself. The experiments that would discriminate among these candidates are the same closed-loop measurements used here, run on a model whose heterogeneity is no longer imposed but generated by cell-autonomous rules, and scored against the same malignant signature. That is the agenda for future work, and the present study supplies both the target it must hit and the negative result that tells it what not to do.

## Conclusions

This study linked the border morphology a clinician observes to the cell-scale rules a modeller can specify, and in doing so it corrected a prevailing expectation, localised a modelling bottleneck, and identified a mechanism that closes the gap. The three contributions move in sequence from measurement to mechanism.

On 1,581 automatically segmented lesions from the ISIC 2018 archive, the malignant signature is not an elevated fractal dimension, as the literature has long assumed, but a specific combination of perimeter- and hull-derived features: compactness rising from 1.351 to 1.810, solidity falling from 0.910 to 0.853, and convexity falling from 0.936 to 0.839.

Together these three describe a border of a few large lobes rather than fine roughness, while the fractal dimension stays nearly constant across grades and is the weakest of the seven descriptors measured, which is itself a useful correction for the field.

The signature survives the most dangerous reading of it, namely that it is a segmentation artefact. The expert ground-truth route reverses five of seven descriptors through a documented polygon-tracing artefact that scales with lesion size, while the automatic route holds all seven in the expected direction and carries roughly an order of magnitude larger effect, and it keeps that separation even when restricted to well-segmented lesions and to the least-hairy subset.

The modelling half of the study is equally instructive. Four canonical mechanisms, including the widely invoked loss of contact inhibition, all converge on near-circular clusters because they share an empty-neighbour division rule that leaves a surrounded cell nowhere to divide. The negative result localises the missing ingredient to that assumption rather than to any single mechanism, which is why it motivates rather than closes the search.

Adding a long-range spatial heterogeneity in the growth rate lifts the model off the circle and reproduces the measured malignant signature to a three-descriptor distance of 0.187, with the correlation length of the field, not its amplitude, identified as the dominant driver by total-effect index in a variance-based sensitivity analysis. The central caveat is also the central contribution: the winning field is imposed rather than emergent, so the study demonstrates sufficiency without yet claiming origin.

Taken together, the results form a single closed loop in which the imaging result names the target, the negative result rules out the obvious route to it, and the alignment result finds a route that works while making its own limitation explicit. That limitation, how a long-range growth-rate heterogeneity could emerge from cell-autonomous dynamics, is the question the study hands forward, posed now with a concrete target to meet and a concrete dead end to avoid. The code, data sources, and configuration are released in a public repository so that the loop can be rerun, extended, and challenged.

## Materials and methods

### Data and lesion grading

The study uses the ISIC 2018 Task 1–2 training archive, comprising dermoscopic images and expert segmentation masks of skin lesions drawn from the International Skin Imaging Collaboration [24]. The challenge archive ships masks but no diagnosis, so a three-tier grade was joined back from the ISIC Archive application programming interface, keyed on the lesion identifier, using collection 63 and the diagnosis_3 field of the hierarchical taxonomy, which maps each lesion to benign nevus, in situ melanoma, or invasive melanoma. The cohort contains 2,223 lesions, of which 1,863 are benign, 198 are in situ melanoma, and 162 are invasive melanoma, and 1,598 carry histopathological confirmation. The images and masks are made available under permissive Creative Commons terms, and the study is secondary use of public, de-identified data, requiring no additional ethics approval beyond the usage terms of each source. The two segmentation routes described below accept overlapping but not identical subsets: the automatic route retains 1,581 lesions (1,348 benign, 124 in situ, 109 invasive) and the expert route retains 2,088 lesions (1,760 benign, 184 in situ, 144 invasive). Lesions are rejected by a common quality-control gate before measurement, described below.

### Segmentation routes and quality control

Every lesion is measured twice, once from the expert ground-truth mask and once from an automatic Otsu segmentation [46], with both routes feeding the identical measurement pipeline so that any disagreement is attributable to segmentation rather than to a second measurement implementation. The automatic route converts the image to the Lab colour space and thresholds the luminance channel with a global Otsu threshold computed only inside the dermatoscope aperture, which prevents the dark vignette corners from biasing the threshold toward the lesion core; on a 90-lesion stratified sample this aperture-cropped threshold alone lifts the mean Dice overlap against the expert masks from 0.554 to 0.632.

Hair suppression uses a DullRazor line structuring element of length one fortieth of the image side in six orientations [47]. Morphological opening is deliberately disabled because it erodes the lesion periphery: with the aperture crop in place, turning opening off raises the recovered area ratio against the expert masks from 0.661 to 0.806. The three settings together yield a fully configured route whose mean Dice overlap reaches 0.672 and whose area ratio reaches 0.822, against 0.644 for the unmodified baseline. The settings were chosen on the 90-lesion sample by minimising the grade-to-grade spread of the area ratio rather than by maximising overlap, because uniform under-segmentation shifts all grades together and largely cancels in a between-grade comparison, whereas grade-dependent under-segmentation would deform the comparison itself in the direction of the hypothesis. The expert route uses the polygon-traced ground-truth masks directly. The automatic route is treated as primary throughout, and the expert route is retained as a sensitivity arm. Masks are never resampled to a common size, since downsampling the largest lesions would erase exactly the fine boundary detail under study and would do so preferentially on the highest-resolution images.

A lesion is rejected before measurement if its equivalent diameter is below 50 pixels, if it occupies more than 80 percent of the image frame, or if more than 2 percent of its boundary lies on the frame edge, the last of which invalidates perimeter and area through truncation. Rejection is deliberately permissive at the frame-contact gate so that large lesions, which run out of frame more often and which correlate with severity, are not differentially discarded; only substantial truncation is excluded.

### Morphological descriptors

Seven boundary shape descriptors are computed on the closed contour extracted from each mask: the box-counting fractal dimension, compactness, circularity, asymmetry, solidity, convexity, and the perimeter decay slope. The fractal dimension is estimated by the box-counting method: the boundary is covered with square grids over 25 logarithmically spaced box sizes, the count of intersecting boxes is recorded at each of nine grid origins to reduce origin bias, and the slope of the least-squares fit of log box-count against log inverse scale yields the fractal dimension. The box sizes are bounded as fractions of the equivalent diameter, from 0.005 to 0.25, with a four-pixel raster floor below which a digital boundary reads as a straight line, so that the fitting window is identical across the roughly tenfold resolution spread of the images. The 0.005 lower bound is chosen because, with typical equivalent diameters of 800 to 2,000 pixels, a coarser ratio would place the lower bound at 16 to 40 pixels and discard the fine-scale detail that distinguishes infiltrative from smooth margins, while also squeezing the fit to roughly 1.1 decades. Rather than maximising the coefficient of determination, the scaling region is detected as a plateau of local slope over a five-point window, with a minimum span of 0.6 decades and at least seven points; a plateau whose slope disagrees with the full-range fit by more than 0.03 is flagged, as is any estimate with a fit coefficient of determination below 0.99 or a slope standard error above 0.03.

Compactness is the squared perimeter divided by four pi times area, solidity is the ratio of area to convex-hull area, convexity is the ratio of perimeter to convex-hull perimeter, circularity is the dispersion of centroid-to-boundary radii, asymmetry is a principal-axis fold overlap of the mask, and the perimeter decay slope is the rate at which the smoothed perimeter shortens as the contour is smoothed at increasing scales. The contour is smoothed with a Gaussian whose width is a fixed fraction of the equivalent diameter, using one twentieth for the reported compactness, so that the correction for raster staircase inflation is resolution-independent.

### Statistical analysis

Between-grade differences are assessed with a Kruskal–Wallis test at a nominal level of 0.05 [48], with the false discovery rate controlled across descriptors by the Benjamini–Hochberg procedure [49] and pairwise comparisons within a descriptor corrected by Holm’s method.

Effect size is reported as eta-squared and as the rank-biserial correlation, and discriminative power is reported as the area under the receiver-operating curve. Confidence intervals are bootstrap percentile intervals from 2,000 resamples with a fixed seed, descriptors with fewer than ten lesions per grade are reported as counts only, and the balanced-subsample sensitivity checks use one hundred replications. Agreement between the two segmentation routes is quantified by the Dice coefficient, and the trend of agreement against severity is tested with the Jonckheere–Terpstra test [50]. All statistical thresholds and random seeds are fixed in a single configuration file so that every constant is traceable.

### Cellular automaton model

The simulation uses a hybrid cellular automaton in the spirit of Anderson’s framework, in which discrete cancer cells occupy an N-by-N lattice while a continuous nutrient field evolves on a coarser grid [30]. A cell becomes able to divide only after reaching a fixed cell-cycle duration, and division requires an empty neighbouring site, so that a cell surrounded by occupied neighbours cannot divide and becomes quiescent; a contact-inhibition factor further gates whether mechanical crowding blocks division. Migration is a random walk to an empty neighbour, biased by a chemotactic term when enabled, and cells die at a small basal rate.

The lattice uses Moore connectivity with a diagonal weight of one over the square root of two to correct grid anisotropy, and the nutrient field obeys a steady-state diffusion–consumption equation expressed in terms of a diffusion length, with far-field, quiescence, and necrosis thresholds on the concentration. The nutrient field is solved on a coarse grid of 128 cells per side, or eight lattice sites per grid cell, and is updated every ten steps; the chemotactic bias is disabled in these experiments, the basal death probability is ten to the minus four, and growth is capped at 6,000 steps although the equivalent-diameter stop usually fires first. The cell-cycle duration is held fixed rather than swept, because it trades against the division probability in a structural non-identifiability that is removed by design and not left to the sensitivity analysis [51].

The mechanism sweep uses a 1,024-by-1,024 lattice and stops growth when the tumour reaches an equivalent diameter of 600 sites, which spans 1.56 decades of box-counting scale and matches the median real lesion. Four candidate mechanisms are swept, each at five levels with three random seeds for 60 runs in total: loss of contact inhibition, in which the contact-inhibition factor is reduced from its strong baseline; clonal heterogeneity, in which a fraction of cells form an advantaged subclone with a tripled division rate; enhanced migration, in which the migration probability and adhesion threshold are varied; and nutrient fingering, in which the diffusion length of the nutrient field is swept from short to long. Every simulated mask is scored by exactly the same measurement code that scored the real lesions, imported by path rather than copied, so that the later alignment is not degraded by a drifted reimplementation. To keep simulated masks comparable to real ones, which carry Otsu segmentation noise, each simulated boundary is also measured after a first-order one-pixel stochastic perturbation of a fixed fraction of its pixels.

### Mechanism alignment and sensitivity analysis

The alignment analysis tests two additional mechanisms at the proliferation rule, on a 512-by-512 lattice grown to an equivalent diameter of 379 sites. The first is mechanical displacement, in which a surrounded cell pushes a neighbouring cell into an empty site, lifting the empty-neighbour bottleneck with an order-one local push. The second is spatial heterogeneity, in which a smooth static growth-rate field multiplies each cell’s division rate; the field has unit mean and is constructed from a small number of low-order Fourier modes, parameterised by an amplitude and a correlation length. Alignment is scored by the distance in the space of the three primary descriptors, compactness, solidity, and convexity, from the simulated boundary to the median invasive-melanoma target, where each descriptor is z-scored by the pooled variation of the real cohort so that the three components are comparable. The coarse sweep compares the two mechanisms at five levels with three seeds. The refinement stage then applies a one-at-a-time screening, a variance-based sensitivity analysis following the Saltelli estimator with a base sample of 256 points and a total of 1,536 evaluations [40–42], and a two-dimensional phase diagram over the amplitude and correlation length at sixteen points per axis. First-order and total-effect indices are computed for the two parameters, and the phase diagram reports the alignment distance across the parameter grid. All measurement, simulation, and alignment code is released in a public repository [52], and every numeric constant described here is declared in a single per-part configuration file so that the pipeline is self-describing and reproducible.

## Acknowledgments

We thank the International Skin Imaging Collaboration for maintaining the ISIC 2018 archive and the ISIC Archive for the diagnostic annotations that made the three-tier grading possible.

## Data and code availability

All analysis code, configuration files, and derived data tables that support the findings of this study are available at https://github.com/neobiosystem/melanoma_morphology_study. The dermoscopic images and expert segmentation masks are publicly available from the International Skin Imaging Collaboration under their original Creative Commons licences, and the diagnostic annotations are available through the ISIC Archive.

## Author contributions

Chang Liu: conceptualization, methodology, software, formal analysis, investigation, data curation, writing (original draft), writing (review and editing), visualization, project administration. Lingling Zu: conceptualization, supervision, writing (review and editing), funding acquisition.

## Supporting information

S1 Fig. Representative dermoscopic lesions with expert segmentations by diagnostic grade. Three example lesions each from the benign, in situ, and invasive grades, with the expert boundary overlaid. Images are from the ISIC 2018 archive and are reproduced under their original CC-0 licences.

## References

1. Arnold M, Singh D, Laversanne M, Vignat J, Vaccarella S, Meheus F, et al. Global burden of cutaneous melanoma in 2020 and projections to 2040. JAMA Dermatology. 2022;158: 495. doi: 10.1001/jamadermatol.2022.0160

2. Bray F, Laversanne M, Sung H, Ferlay J, Siegel RL, Soerjomataram I, et al. Global cancer statistics 2022: GLOBOCAN estimates of incidence and mortality worldwide for 36 cancers in 185 countries. CA: A Cancer Journal for Clinicians. 2024;74: 229–263. doi: 10.3322/caac.21834

3. Friedman RJ, Rigel DS, Kopf AW. Early detection of malignant melanoma: The role of physician examination and self-examination of the skin. CA: A Cancer Journal for Clinicians. 1985;35: 130–151. doi: 10.3322/canjclin.35.3.130

4. Abbasi NR, Shaw HM, Rigel DS, Friedman RJ, McCarthy WH, Osman I, et al. Early diagnosis of cutaneous melanoma: Revisiting the ABCD criteria. JAMA. 2004;292: 2771–2776. doi: 10.1001/jama.292.22.2771

5. Friedl P, Alexander S. Cancer invasion and the microenvironment: Plasticity and reciprocity. Cell. 2011;147: 992–1009. doi: 10.1016/j.cell.2011.11.016

6. Claridge E, Hall PN, Keefe M, Allen JP. Shape analysis for classification of malignant melanoma. Journal of Biomedical Engineering. 1992;14: 229–234. doi: 10.1016/0141-5425(92)90057-R

7. Carbonetto SH, Lew SE. Characterization of border structure using fractal dimension in melanomas. Annual international conference of the IEEE engineering in medicine and biology society (EMBC). 2010. doi: 10.1109/IEMBS.2010.5627296

8. Styła M, Giżewski T. The study of usefulness of a set of fractal parameters to build classes of disease units based on images of pigmented skin lesions. Diagnostics. 2021;11: 1773. doi: 10.3390/diagnostics11101773

9. Celebi ME, Kingravi HA, Iyatomi H, Aslandogan YA, Stoecker WV, Moss RH, et al. Border detection in dermoscopy images using statistical region merging. Skin Research and Technology. 2008;14: 347–353. doi: 10.1111/j.1600-0846.2008.00301.x

10. Mirikharaji Z, Abhishek K, Bissoto A, Barata C, Avila S, Valle E, et al. A survey on deep learning for skin lesion segmentation. Medical Image Analysis. 2023;88: 102863. doi: 10.1016/j.media.2023.102863

11. Dong Y, Wang L, Li Y. TC-net: Dual coding network of transformer and CNN for skin lesion segmentation. PLOS ONE. 2022;17: e0277578. doi: 10.1371/journal.pone.0277578

12. Amin J, Azhar M, Arshad H, Zafar A, Kim S-H. Skin-lesion segmentation using boundary-aware segmentation network and classification based on a mixture of convolutional and transformer neural networks. Frontiers in Medicine. 2025;12. doi: 10.3389/fmed.2025.1524146

13. Ali A-R, Li J, Yang G, O’Shea SJ. A machine learning approach to automatic detection of irregularity in skin lesion border using dermoscopic images. PeerJ Computer Science. 2020;6: e268. doi: 10.7717/peerj-cs.268

14. Esteva A, Kuprel B, Novoa RA, Ko J, Swetter SM, Blau HM, et al. Dermatologist-level classification of skin cancer with deep neural networks. Nature. 2017;542: 115–118. doi: 10.1038/nature21056

15. Brinker TJ, Hekler A, Enk AH, Klode J, Hauschild A, Berking C, et al. Deep learning outperformed 136 of 157 dermatologists in a head-to-head dermoscopic melanoma image classification task. European Journal of Cancer. 2019;113: 47–54. doi: 10.1016/j.ejca.2019.04.001

16. Tschandl P, Rinner C, Apalla Z, Argenziano G, Codella N, Halpern A, et al. Human–computer collaboration for skin cancer recognition. Nature Medicine. 2020;26: 1229–1234. doi: 10.1038/s41591-020-0942-0

17. Haggenmüller S, Maron RC, Hekler A, Utikal JS, Barata C, Barnhill RL, et al. Skin cancer classification via convolutional neural networks: Systematic review of studies involving human experts. European Journal of Cancer. 2021;156: 202–216. doi: 10.1016/j.ejca.2021.06.049

18. Patel RH, Foltz EA, Witkowski A, Ludzik J. Analysis of artificial intelligence-based approaches applied to non-invasive imaging for early detection of melanoma: A systematic review. Cancers. 2023;15: 4694. doi: 10.3390/cancers15194694

19. Daneshjou R, Vodrahalli K, Novoa RA, Jenkins M, Liang W, Rotemberg V, et al. Disparities in dermatology AI performance on a diverse, curated clinical image set. Science Advances. 2022;8. doi: 10.1126/sciadv.abq6147

20. Ahmad M, Ahmed I, Chehri A, Jeon G. Fusion of metadata and dermoscopic images for melanoma detection: Deep learning and feature importance analysis. Information Fusion. 2025;124: 103304. doi: 10.1016/j.inffus.2025.103304

21. Naseri H, Safaei AA. Diagnosis and prognosis of melanoma from dermoscopy images using machine learning and deep learning: A systematic literature review. BMC Cancer. 2025;25. doi: 10.1186/s12885-024-13423-y

22. Tschandl P, Rosendahl C, Kittler H. The HAM10000 dataset, a large collection of multi-source dermatoscopic images of common pigmented skin lesions. Scientific Data. 2018;5: 180161. doi: 10.1038/sdata.2018.161

23. Codella NCF, Gutman D, Celebi ME, Helba B, Marchetti MA, Dusza SW, et al. Skin lesion analysis toward melanoma detection: A challenge at the 2017 international symposium on biomedical imaging (ISBI), hosted by the international skin imaging collaboration (ISIC). 2018 IEEE 15th International Symposium on Biomedical Imaging (ISBI 2018). 2018; 168–172. doi: 10.1109/isbi.2018.8363547

24. Codella NCF, Rotemberg V, Tschandl P, Celebi ME, Dusza S, Gutman D, et al. Skin lesion analysis toward melanoma detection 2018: A challenge hosted by the international skin imaging collaboration (ISIC). 2019. Available: https://arxiv.org/abs/1902.03368

25. Rotemberg V, Kurtansky N, Betz-Stablein B, Caffery L, Chousakos E, Codella N, et al. A patient-centric dataset of images and metadata for identifying melanomas using clinical context. Scientific Data. 2021;8. doi: 10.1038/s41597-021-00815-z

26. Altrock PM, Liu LL, Michor F. The mathematics of cancer: Integrating quantitative models. Nature Reviews Cancer. 2015;15: 730–745. doi: 10.1038/nrc4029

27. Anderson ARA, Quaranta V. Integrative mathematical oncology. Nature Reviews Cancer. 2008;8: 227–234. doi: 10.1038/nrc2329

28. Macnamara CK. Biomechanical modelling of cancer: Agent-based force-based models of solid tumours within the context of the tumour microenvironment. Computational and Systems Oncology. 2021;1. doi: 10.1002/cso2.1018

29. Bull JA, Byrne HM. Quantification of spatial and phenotypic heterogeneity in an agent-based model of tumour-macrophage interactions. PLOS Computational Biology. 2023;19: e1010994. doi: 10.1371/journal.pcbi.1010994

30. Anderson ARA. A hybrid mathematical model of solid tumour invasion: The importance of cell adhesion. Mathematical Medicine and Biology. 2005;22: 163–186. doi: 10.1093/imammb/dqi005

31. Cheung KJ, Horne-Badovinac S. Collective cell migration modes in development, tissue repair and cancer. Nature Reviews Molecular Cell Biology. 2025;26: 741–758. doi: 10.1038/s41580-025-00858-9

32. Xu X, Xu J, Liu J, Jiang C, Tian L, Xu Y, et al. Cellular mechanics of finger-like structures of collective cell migration. Journal of Applied Mechanics. 2023;91. doi: 10.1115/1.4063217

33. Hanahan D, Weinberg RA. Hallmarks of cancer: The next generation. Cell. 2011;144: 646–674. doi: 10.1016/j.cell.2011.02.013

34. Drasdo D, Höhme S. A single-cell-based model of tumor growth in vitro: Monolayers and spheroids. Physical Biology. 2005;2: 133–147. doi: 10.1088/1478-3975/2/3/001

35. Byrne H, Drasdo D. Individual-based and continuum models of growing cell populations: A comparison. Journal of Mathematical Biology. 2008;58: 657–687. doi: 10.1007/s00285-008-0212-0

36. Wang J, Zhang L, Jing C, Ye G, Wu H. Multi-scale agent-based modeling on melanoma and its related angiogenesis analysis. Theoretical Biology and Medical Modelling. 2013;10: 41. doi: 10.1186/1742-4682-10-41

37. Brú A, Pastor JM, Fernaud I, Brú I, Melle S. Super-rough dynamics on tumor growth. Physical Review Letters. 1998;81: 4008–4011. doi: 10.1103/PhysRevLett.81.4008

38. van der Laak J, Litjens G, Ciompi F. Deep learning in histopathology: The path to the clinic. Nature Medicine. 2021;27: 775–784. doi: 10.1038/s41591-021-01343-4

39. Baxi V, Edwards R, Montalto M, Saha S. Digital pathology and artificial intelligence in translational medicine and clinical practice. Modern Pathology. 2022;35: 23–32. doi: 10.1038/s41379-021-00919-2

40. Homma T, Saltelli A. Importance measures in global sensitivity analysis of nonlinear models. Reliability Engineering & System Safety. 1996;52: 1–17. doi: 10.1016/0951-8320(96)00002-6

41. Saltelli A, Annoni P, Azzini I, Campolongo F, Ratto M, Tarantola S. Variance based sensitivity analysis of model output. Design and estimator for the total sensitivity index. Computer Physics Communications. 2010;181: 259–270. doi: 10.1016/j.cpc.2009.09.018

42. Razavi S, Jakeman A, Saltelli A, Prieur C, Iooss B, Borgonovo E, et al. The future of sensitivity analysis: An essential discipline for systems modeling and policy support. Environmental Modelling & Software. 2021;137: 104954. doi: 10.1016/j.envsoft.2020.104954

43. Gerlee P, Anderson ARA. The evolution of carrying capacity in constrained and expanding tumour cell populations. Physical Biology. 2015;12: 056001. doi: 10.1088/1478-3975/12/5/056001

44. Black JRM, McGranahan N. Genetic and non-genetic clonal diversity in cancer evolution. Nature Reviews Cancer. 2021;21: 379–392. doi: 10.1038/s41568-021-00336-2

45. Turing AM. The chemical basis of morphogenesis. Philosophical Transactions of the Royal Society of London Series B, Biological Sciences. 1952;237: 37–72. doi: 10.1098/rstb.1952.0012

46. Otsu N. A threshold selection method from gray-level histograms. IEEE Transactions on Systems, Man, and Cybernetics. 1979;9: 62–66. doi: 10.1109/tsmc.1979.4310076

47. Lee T, Ng V, Gallagher R, Coldman A, McLean D. Dullrazor®: A software approach to hair removal from images. Computers in Biology and Medicine. 1997;27: 533–543. doi: 10.1016/s0010-4825(97)00020-6

48. Kruskal WH, Wallis WA. Use of ranks in one-criterion variance analysis. Journal of the American Statistical Association. 1952;47: 583–621. doi: 10.1080/01621459.1952.10483441

49. Benjamini Y, Hochberg Y. Controlling the false discovery rate: A practical and powerful approach to multiple testing. Journal of the Royal Statistical Society Series B: Statistical Methodology. 1995;57: 289–300. doi: 10.1111/j.2517-6161.1995.tb02031.x

50. Jonckheere AR. A distribution-free k-sample test against ordered alternatives. Biometrika. 1954;41: 133–145. doi: 10.2307/2333011

51. Simpson MJ, Baker RE. Parameter identifiability, parameter estimation, and model prediction for differential equation models. SIAM Review. 2026;68: 153–171. doi: 10.1137/24m1667968

52. Cadwallader L, Mac Gabhann F, Papin J, Pitzer VE. Advancing code sharing in the computational biology community. PLOS Computational Biology. 2022;18: e1010193. doi: 10.1371/journal.pcbi.1010193

